# Recapitulating the notochord to nucleus pulposus transition from human mesendoderm progenitors in 3D

**DOI:** 10.64898/2026.10.05.756547

**Authors:** Julie Warin, Nicolas Vedrenne, Bluwen Guidoux d’Halluin, Claire Chédeville, Maëlle Blanche, Clémence Maillard, Caroline Chariau, Christabel Dube, Romain Desprat, Benjamin Gantenbein, Stephen M. Richardson, Jérôme Guicheux, Laurent David, Marianna A. Tryfonidou, Jean-Marc Lemaître, Anne Camus

## Abstract

Intervertebral disc (IVD) degeneration, a major cause of low back pain, is driven by the disruption of nucleus pulposus (NP) tissue homeostasis. The NP arises from embryonic notochord cells that differentiate and adapt to the hypoxic, matrix-rich environment of the developing disc. Recapitulating the developmental trajectory from notochordal progenitors to NP-like cells *in vitro* remains challenging, as most protocols bypass early specification steps and fail to capture this understudied lineage. Here, using a three-dimensional (3D) system, we reconstruct this developmental path from human pluripotent stem cells into mesendoderm progenitor cells (MEPCs), notochord-like cells (NLCs), and NP-like cells, guided by single-cell transcriptomics and spatial profiling. We identify MEPCs as a distinct progenitor population with a unique gene signature and conserved regulatory networks (*GSC*, *KLF8*, *LHX1*, *MIXL1*) shared with early organizer and mesendodermal cells. Self-organization in MEPCs-derived aggregates induced spontaneous expression of *NOTO*, a key transcription factor orchestrating notochordal identity, and enhanced commitment to notochordal fate. Long-term culture in a medium containing TGF-β1 and GDF-5 promoted maturation into SOX9^+^, KRT8^+^, CD109^+^ NP-like cells exhibiting robust matrix deposition and prominent LAMP1-positive intracytoplasmic vacuoles. This scalable, physiologically relevant model enables fundamental studies of human disc biology and early degenerative processes, while advancing regenerative strategies.

## Introduction

During mammalian embryogenesis, nucleus pulposus cells originate from the notochord, a transient axial structure that regulates vertebral patterning, serves as a critical signaling center for proper axial skeleton development, and ultimately contributes to intervertebral disc (IVD) formation^1^. Notably, notochord-derived cells persist in the core of the developing IVD, where they form the nucleus pulposus (NP)^2,3^, the proteoglycan-rich and highly hydrated tissue compartment. The NP is often the primary site of observed degenerative changes, including marked inflammation mediating extracellular matrix (ECM) breakdown, loss of hydration, drastic cellular phenotypic changes, and decreased cellularity^4^. This cascade of structural and functional alterations contributes to IVD degeneration and is strongly associated with low back pain^5^ and the decline of the notochordal cells (NCs)^6^.

NP developmental origin is well established and the transition from the notochord to NP cells largely described during disc development and growth in animal models^6–8^. However, the progressive differentiation of the NCs into NP tissue, as well as the mechanisms that regulate this process, remain poorly characterized^2,3^ and a significant challenge in humans^1^. Indeed, species-specific variations in NP composition and NCs persistence limit their translational relevance^6^. In humans, NCs disappear before birth, and their complete loss by skeletal maturity is associated with disc degeneration. In contrast, the disc remains protected in species, such as mice or non-chondrodystrophic dogs, where NCs persist throughout life underscoring the regenerative potential of the NCs population^9^. Human NP single-cell transcriptomic atlases have been constructed using samples collected during embryonic and postnatal development, as well as at different stages of life and varying grades of degeneration. They reveal the wide cellular heterogeneity and have enabled the identification of distinct subpopulations of NP cells, including progenitors and mature cells, as well as cells related to the degenerative process, including fibrocartilage-associated cells, and stress- or inflammation-associated cells^10–16^.

The healthy NP contains two well-described and distinct cell populations: large, vacuolated NCs, prominent in early life, and smaller non-vacuolated nucleopulpocytes (NPCs), predominant in adulthood and sharing matrix and molecular features with articular cartilage chondrocytes^17^. The replacement of NCs by NPCs is a hallmark of disc aging and is widely accepted as reflecting NCs maturation and terminal differentiation into NPCs. Physicochemical cues, such as hypoxia, osmotic and mechanical stress, and nutrient limitation, contribute to this phenotypic shift^18–26^. Progressively, vacuolated progenitor NCs give place to NPCs that exhibit significantly higher expression of Aggrecan, Collagen II, and Sox9 transcription factor, which are essential for ECM homeostasis and the biomechanical properties of the disc^22^. Conversely, this results in a marked reduction in NCs paracrine activity, which numerous *in vitro* studies have shown supports NPCs survival and function^20,25,27–30^, and inhibits neo-innervation and neo-vascularization within the IVD, two processes associated with degeneration^31,32^. The presence of NCs correlates with a healthier, regenerative disc phenotype. Thus, deeper insights into their phenotypic transition and the mechanisms governing NC-to-NPC crosstalk in the human NP are critical for elucidating the causal onset and progression of degenerative disc disease^6,33^. Even more so, in a therapeutic context of degenerative disc disease^31^, NCs represent a key target for innovative treatment strategies^6^ owing to their central role in NP formation and disc homeostasis, as well as their potential to repair degenerative discs. However, studying these processes in human tissues remains problematic due to the limited availability of NC-rich NPs and the inability to expand primary NCs *in vitro*^34^, making induced pluripotent stem cells (iPSC)-based models essential for both mechanistic investigations and future translational applications.

Human iPSC-based models have been developed to generate notochordal-like cells (NLCs). They often rely on the expression of the notochord-related transcription *NOTO* and typically yield immature embryonic notochordal states, associated with a non-vacuolated phenotype and poor matrix synthesis^11,35–37^. Despite extended culture to promote NLCs differentiation toward glycosaminoglycans (GAGs)-rich matrix-producing cells, the resulting cell identity often remains insufficiently resolved due to reliance on a limited set of NP markers^38–40^. Current differentiation protocols often bypass early developmental stages, limiting our understanding of the transcriptional programs that guide notochord specification and NP cell formation. More advanced 3D systems, including gastruloids and segmentoids have provided valuable insights into early embryonic events, but they only capture the initial stage of notochordal lineage specification or fail to form a *bona fide* notochord, limiting their relevance for modeling IVD development^41–48^. Finally, other common approaches used adult mesenchymal or adipose stem cells for their chondrogenic potency. These models, however, suffer from patient-to-patient variability and lack developmental relevance to the notochordal lineage^49–51^. Yet, the lack of consensus on effective growth factor combinations, coupled with the reliance broad chondrogenic markers such as Collagen II and Aggrecan to infer NP-like identity hinders cross study comparisons^32^.Collectively, these limitations underscore the need for standardized *in vitro* models that recapitulate the developmental continuum from progenitors to juvenile human NP tissue and corresponding markers to accurately define NP cell identity. These gaps represent a major challenge that impedes both the understanding human development and the advancement of effective regenerative therapies.

Our study addresses part of this challenge by establishing and validating a 3D *in vitro* model based upon self-aggregation of transcriptionally defined human iPSCs-derived progenitors to recapitulate features of NP development. Exposure to controlled growth factors, including TGFβ1 and GDF-5, over 28 days of culture directed sequential differentiation into NLCs and ultimately into functional NP-like cells producing matrix. Time-course transcriptomic profiling, qualitative and quantitative ECM analysis, immunohistochemistry, and spatial gene expression mapping confirmed the progressive acquisition of NP identity and indicated the coexistence of the two major cell types found in the native juvenile human disc. Overall, this hiPSC-based 3D culture system provides a developmentally grounded and physiologically relevant model to investigate notochord and NP biology, offering first insights into key mechanisms that may be exploited to regenerative approaches.

## Results

### Human iPSC-derived mesendoderm progenitors display a unique molecular signature and organizer-like transcriptional programs

Previous studies have evaluated the requirements of Wnt and Nodal signaling pathways for the generation of mesendoderm progenitor cells (MEPCs)^36,37,52^. These early developmental progenitors co-express the transcription factors *TBXT* and *FOXA2* and, *in vitro,* have the potential to differentiate into endoderm and mesoderm derivatives^11^. In order to define the molecular signature from MEPCs to NLCs, single-cell RNA sequencing (scRNAseq) was performed to profile differentiating hiPSCs from day 2 to day 7 (Fig. 1a, b). Unsupervised clustering revealed 13 distinct cell clusters, each corresponding to differentiating populations at successive developmental stages (Fig. 1c, Fig. S1a, Table S1). MEPCs were exclusively detected at day 2. Clustering revealed day-3 commitment toward endodermal and mesodermal fates, consistent with progenitor-like behavior (Fig. 1b,c, Fig. S1a). From day 5 to day 7, differentiated endodermal and mesodermal derivatives emerged, together with a distinct notochord cluster previously identified as highly similar to the human fetal notochord^11^ (Fig. 1b,c, Fig. S1 a,b).

**Figure 1.**
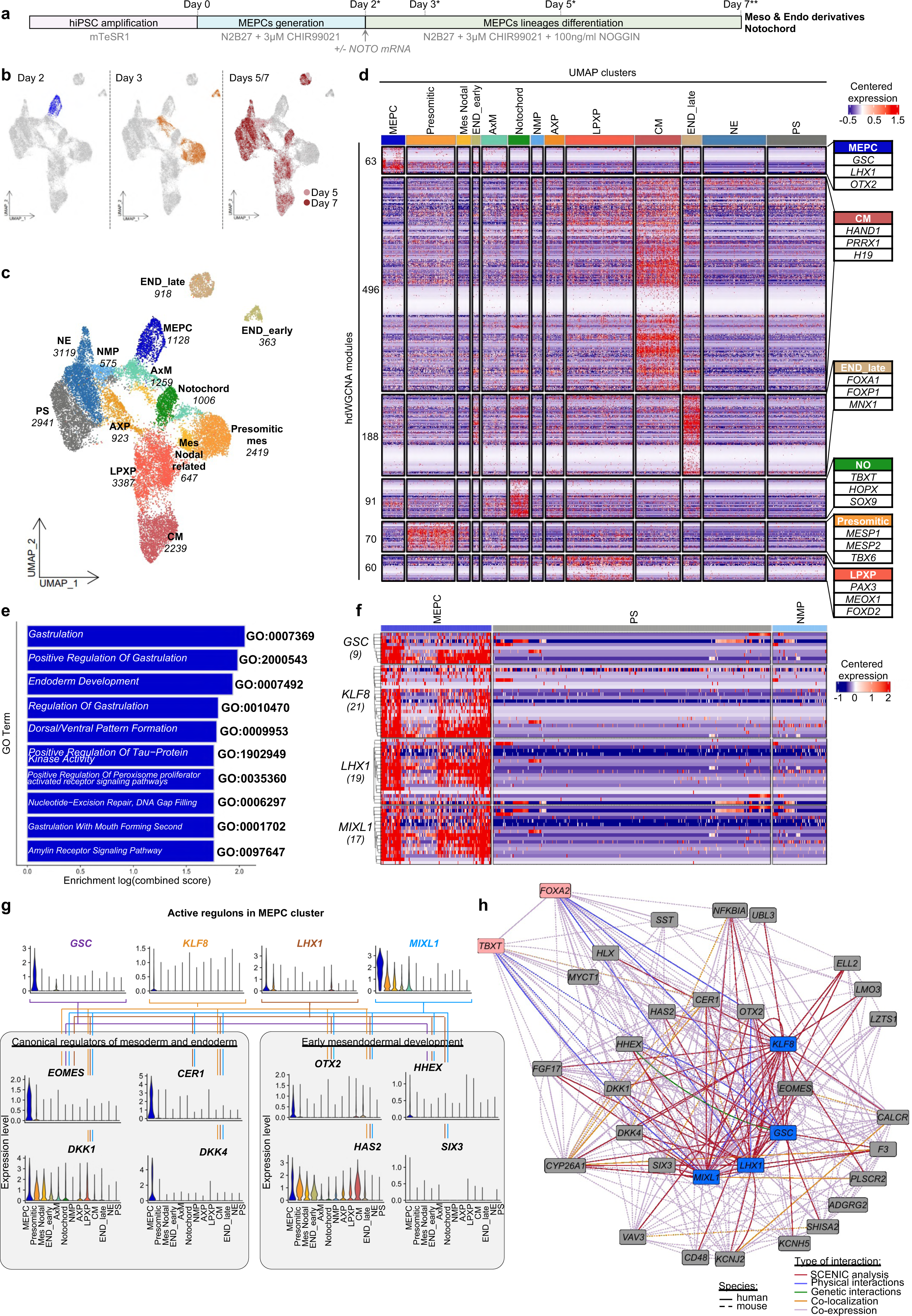
Molecular characterization of mesendoderm progenitors at the single-cell level. (a) Overview of the differentiation protocol and scRNA-seq strategy. *indicates scRNAseq time points generated in this study, **indicates scRNAseq time points generated in Warin et al. 2024^11^ (b) UMAP projection of single-cell transcriptomes colored by time point, showing progression over differentiation. Days 5 and 7 conditions are projected together as they cluster together. (c) UMAP visualization of unsupervised clustering following batch correction, identifying 13 lineage-associated clusters. The number of cells in each cluster is indicated. (d) Heatmap revealing gene modules across cell clusters following Weighted Gene Co-expression Network Analysis (WGCNA). Each row represents a module of co-expressed genes; row height corresponds to the number of genes per module, and gene numbers are indicated. Columns represent individual cells. Gene expression levels are color-coded from low (blue) to high (red). Only modules relevant to the annotated clusters are shown. (e) Gene Ontology term enrichment for the gene module associated with the MEPCs. Bars represent significantly enriched biological processes; GO term IDs are listed on the right. Bar length reflects enrichment significance. (f) Heatmap showing expression of SCENIC-inferred regulons within the MEPCs at day 2. Only clusters with more than 18 cells at this time point are displayed. Gene expression of MEPCs-specific regulons in the primitive streak (PS) and neuromesodermal progenitors (NMP) is shown for comparison. Rows represent individual regulons, with expression levels color-coded from low (blue) to high (red). The number of targets is indicated in brackets. (g) Network visualization of selected SCENIC regulons with key target genes. Violin plots indicate the expression levels of selected regulons and their target genes across annotated clusters. (h) Functional gene interaction network generated on the SCENIC-inferred target genes using GeneMANIA. Nodes represent query genes, and edge colors indicate interaction types. Only interactions with weights > 0.1 are shown. TBXT and FOXA2 are widely expressed in the dataset, and are not specific to the MEPCs cluster. Because to their relevance, we manually added them for the GeneMania analysis (red boxes). Regulons are highlighted in blue boxes. Abbreviations: AxM: Axial mesoderm; MEPC: mesendoderm progenitors; END: endoderm; mes: mesoderm; CM: cardiac mesoderm; PS: primitive streak; NE: neuroectoderm; NMP: neuromesodermal progenitors; AXP: axial progenitors; LPXP: lateral/paraxial progenitors; NO: notochord.

To gain insights into the molecular program defining specific cell populations, the expression of correlated gene modules was analyzed across all clusters (Fig. 1d). Beyond the previously described notochord-associated module enriched for markers such as *TBXT*, *HOPX,* and *SOX9*^11^ ([Gitlink](https://github.com/w-julie/3D-cell-paper/blob/main/scRNAseq/2-hdWGCNA/Modules-genelists/pink-notochord.csv)), this analysis identified distinct transcriptional modules and in particular a module comprising 63 genes uniquely associated with the MEPC cluster ([Gitlink](https://github.com/w-julie/3D-cell-paper/blob/main/scRNAseq/2-hdWGCNA/Modules-genelists/greenyellow-MEPC.csv)). Gene Ontology (GO) analysis linked the MEPC-specific module to biological processes related to gastrulation (GO:0007369, GO:2000543, GO:0010470, GO:0001702), endoderm development (GO:0007492), and dorsal/ventral pattern formation (GO:0009953) (Fig. 1e). Notably, the murine mesendoderm progenitor’s gene signature from Masamsetti et al., 2025^53^ was selectively enriched in the hiPSC-derived MEPC cluster (Fig. S1c).

To further characterize the regulatory mechanisms underlying MEPCs identity and developmental potential, transcription factor activity was inferred using SCENIC, identifying four active regulons, *GSC, KLF8, LHX1* and *MIXL1,* associated with organizer and axial mesendoderm-related populations *in vivo*^53–55^ (Fig. 1f). These regulons’ predicted target genes were selectively expressed in MEPCs and were largely absent from the primitive streak (PS) and neuromesodermal progenitor (NMP) clusters (Fig. 1f, [Gitlink](https://github.com/w-julie/3D-cell-paper/tree/main/scRNAseq/3-SCENIC/Gene-list-regulons-filtered-hdWGCNA)). SCENIC-inferred target genes revealed several canonical regulators of mesoderm and endoderm development, including *EOMES*, *CER1*, *DKK1* and *DKK4* (Fig. 1g), with *CER1* and *DKK1* specifically reported to be expressed in the mouse mesendoderm progenitors at embryonic days 7-7.5^53,56,57^ and in the human axial mesoderm cells at Carnegie Stage 6-8^10,58^. Additional SCENIC-inferred targets, including *OTX2, HHEX, HAS2,* and *SIX3*, further supported the relevance of the identified regulatory networks to early mesendoderm development. Gene network analysis highlighted the regulatory architecture underlying MEPC-associated transcriptional programs, including highly connected genes and predicted functional associations within the MEPC module (Fig. 1h). Furthermore, the data highlighted physical interactions between *LHX1, OTX2* and *FOXA2 genes*^55^, as well as genetic interactions between *GSC* and *HHEX*^59^ reported in mouse and Xenopus respectively.

### Three-dimensional self-organization promotes early notochordal specification of mesendoderm progenitors

To determine whether 3D culture is sufficient to support notochordal commitment independently of exogenous *NOTO* mRNA^11,37^, MEPCs were dissociated into single cells, transfected or not with *NOTO* mRNA, allowed to self-aggregate under low-attachment conditions, and cultured for an additional 5 days (Fig. 2a). The spatial organization of NLCs within the aggregates were assessed by immunofluorescence at day 7. TBXT/FOXA2 and TBXT/SOX9 co-stainings confirmed the presence of NLCs in both conditions (Fig. 2b). NLCs were preferentially localized to internal regions of the aggregates, suggesting a permissive local microenvironment. A time-course analysis of NLCs marker expression was performed to compare differentiation dynamics between 3D and standard 2D monolayer cultures in matched medium composition (Fig. 2c). On days 3 and 7, endogenous *NOTO* expression was significantly higher in both transfected and not transfected aggregates than in monolayer cultures. Consistent with the defined genetic hierarchy involving *NOTO* and governing node and notochord development in the mouse embryo^60,61^, *TBXT* and *FOXA2* transcripts showed higher expression on day 3 in aggregates (Fig. 2c). *SOX9* expression increased over time and, remarkably, by day 7 reached comparable levels in transfected and not transfected aggregates (Fig. 2c). Despite elevated *NOTO* expression, the notochord-specific morphogen SHH^62^ showed only slight upregulation in aggregates (less than *NOTO*, though *SHH* transcripts exceeded hiPSCs levels) indicating further maturation is needed.

**Figure 2.**
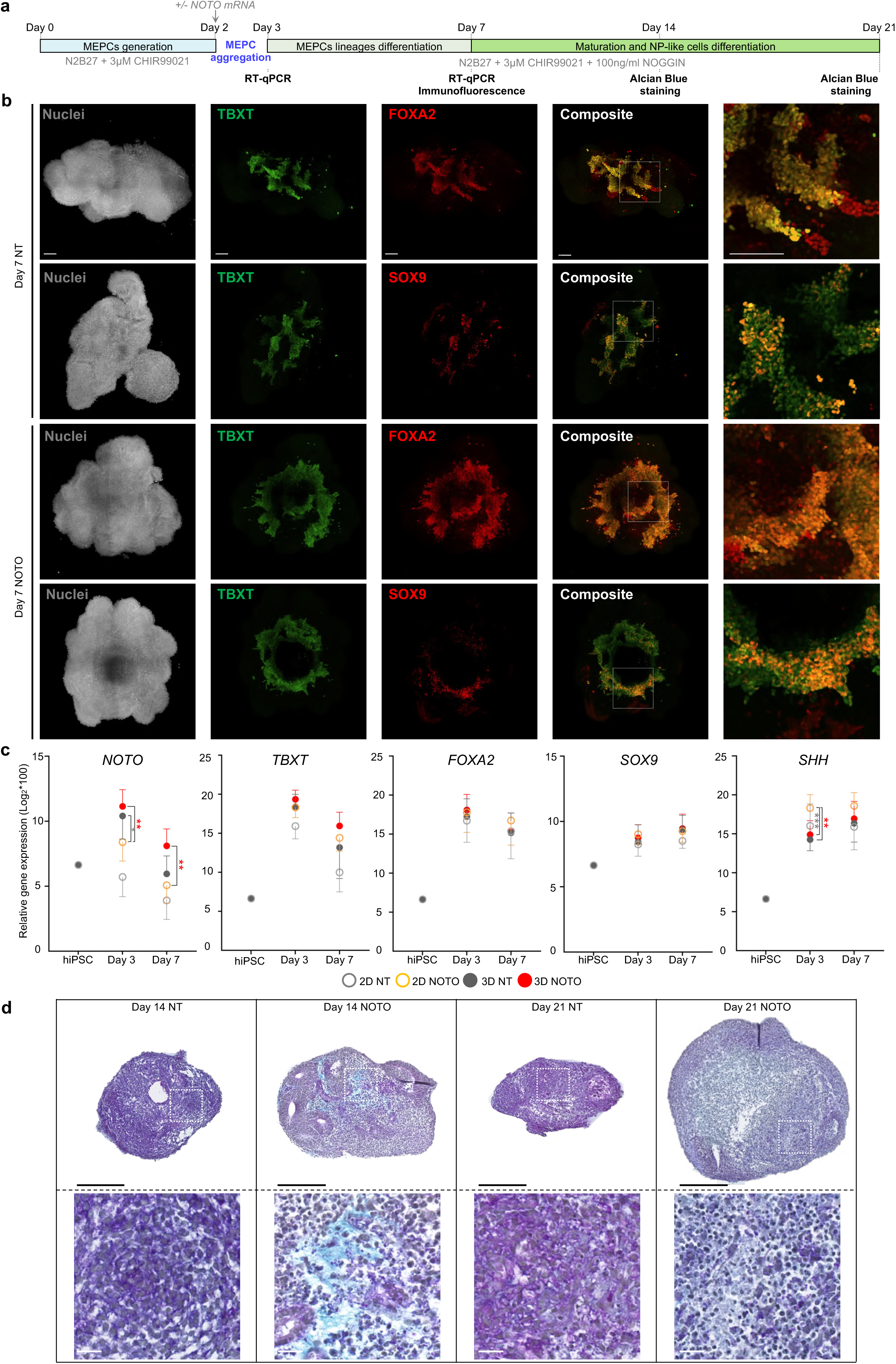
Self-organization drives MEPCs differentiation into NLCs. (a) Schematic representation of the 3D culture system used to promote MEPCs self-organization and differentiation. (b) 3D projection of representative co-immunostainings for TBXT/FOXA2 and TBXT/SOX9 at day 7, on 3D-aggregates of both *NOTO*-transfected and non-transfected conditions. High-magnification imaging further revealed the presence of cells co-expressing or expressing TBXT, FOXA2 or SOX9 individually in both transfected and non-transfected aggregates. Scale bars, 100 µm; insets, 50 µm. (c) RT-qPCR analysis of key notochordal markers (NOTO, TBXT, FOXA2, SOX9, and SHH) at day 3 (i.e., 24h after transfection and aggregation) and day 7 relative to unstimulated hiPSCs. Data represent fold-change (mean ± s.e.m.) from 6-8 biological replicates across four independent hiPSC lines. Statistical comparisons between monolayer NOTO-transfected and 3D NT/NOTO conditions were performed using the Mann-Whitney test (*p < 0.05; **p < 0.001). (d) Alcian blue staining of 3D aggregates at day 14 and day 21. Nuclei are visualized in blue-purple and glycosaminoglycans (GAGs) in blue. Scale bars: top panel, 250μm; bottom panel, 25μm.

To further assess NLCs maturation, structural organization and matrix production, aggregates were analysed over time in long-term culture. At days 14 and 21, *NOTO*-transfected aggregates showed modest growth compared to not transfected aggregates that remained compact (Fig. 2d). Cells exhibited neither vacuole-like structures nor the rounded morphology characteristic of NPC-like cells (Fig. 2d), indicating an immature state. Minimal glycosaminoglycans (GAGs) deposition was detected, exclusively in the *NOTO*-transfected conditions, in spite of attempts to stimulate ECM production using ascorbic acid and Insulin-transferrin-sodium selenite (ITS) supplementation, known to enhance collagen and GAG synthesis^63^ (data not shown). Collectively, these findings highlight the 3D culture environment is sufficient to increase endogenous *NOTO* expression and early notochordal markers. The 3D physical constraints alone were insufficient to support further maturation suggesting that additional developmental cues may be necessary for full notochordal maturation.

### TGF-β and GDF signalling induce matrix-producing differentiation and phenotypic maturation in 3D culture

To address limited notochordal maturation observed under 3D self-organization alone, we evaluated differentiation strategies based on developmental cues known to enhance matrix-producing phenotypes in stem cells^32^. Aggregates were cultured for up to 28 days under hypoxic conditions (5% O2^50^) and exposed to three different media formulations all supplemented with ascorbic acid, dexamethasone, and ITS: (i) chondrogenic differentiation medium supplemented with TGF-β1 (CHM), which is reported to induce chondrocyte differentiation programs^49^, (ii) nucleopulpogenic differentiation medium supplemented with TGF-β1 and GDF5 (NPM), which more specifically promotes NP-related matrix synthesis^49^, and (iii) a sequential exposure to CHM and NPM differentiation media (CHM-NPM), which enhances ECM production^36^. To evaluate maturation capacity under 3D culture conditions, aggregates derived from NLCs were compared with aggregates derived from the earlier progenitor population MEPCs. Following their generation in monolayer culture and validation of differentiation efficiency (Fig. S2), both cell populations were dissociated, aggregated, and cultured in 3D for extended periods before comparative analysis (Fig. 3a). NLCs-derived aggregates growth was observed over time, as measured by the area of the aggregates, with comparable sizes detected in CHM and NPM conditions, whereas aggregates cultured under CHM-NPM condition were approximately 1.6-fold larger by day 28 (Fig. 3b). Aggregates growth was associated with increased deposition of sulfated GAGs, as confirmed by quantitative assays (Fig. 3c, d). Over time, Alcian Blue staining revealed two distinct cellular organizations within the aggregates: one characterized by abundant ECM deposition and the other by densely packed cells (Fig. 3e). MEPCs-derived aggregates exhibited progressive growth up to day 28, with larger sizes under NPM and CHM-NPM conditions than under CHM (Fig. 3f). As with NLCs-aggregates, ECM production began by day 7 and increased over time. At day 28, significantly higher amounts of sulfated GAGs were deposited by MEPCs-derived aggregates in NPM and CHM-NPM than in CHM (p=0.0079, Fig. 3g). Remarkably, MEPCs_NPM aggregates formed a homogeneous, ECM-enriched cellular organization, while aggregates under CHM and CHM-NPM exhibited a distinct ECM-rich organization, separate from their densely cellular domains (Fig. 3h, i). By day 28, prominent vacuole-like structures were detected in MEPCs_NPM aggregates (arrows in Fig. 3i), consistent with progression toward a more mature notochordal phenotype.

**Figure 3.**
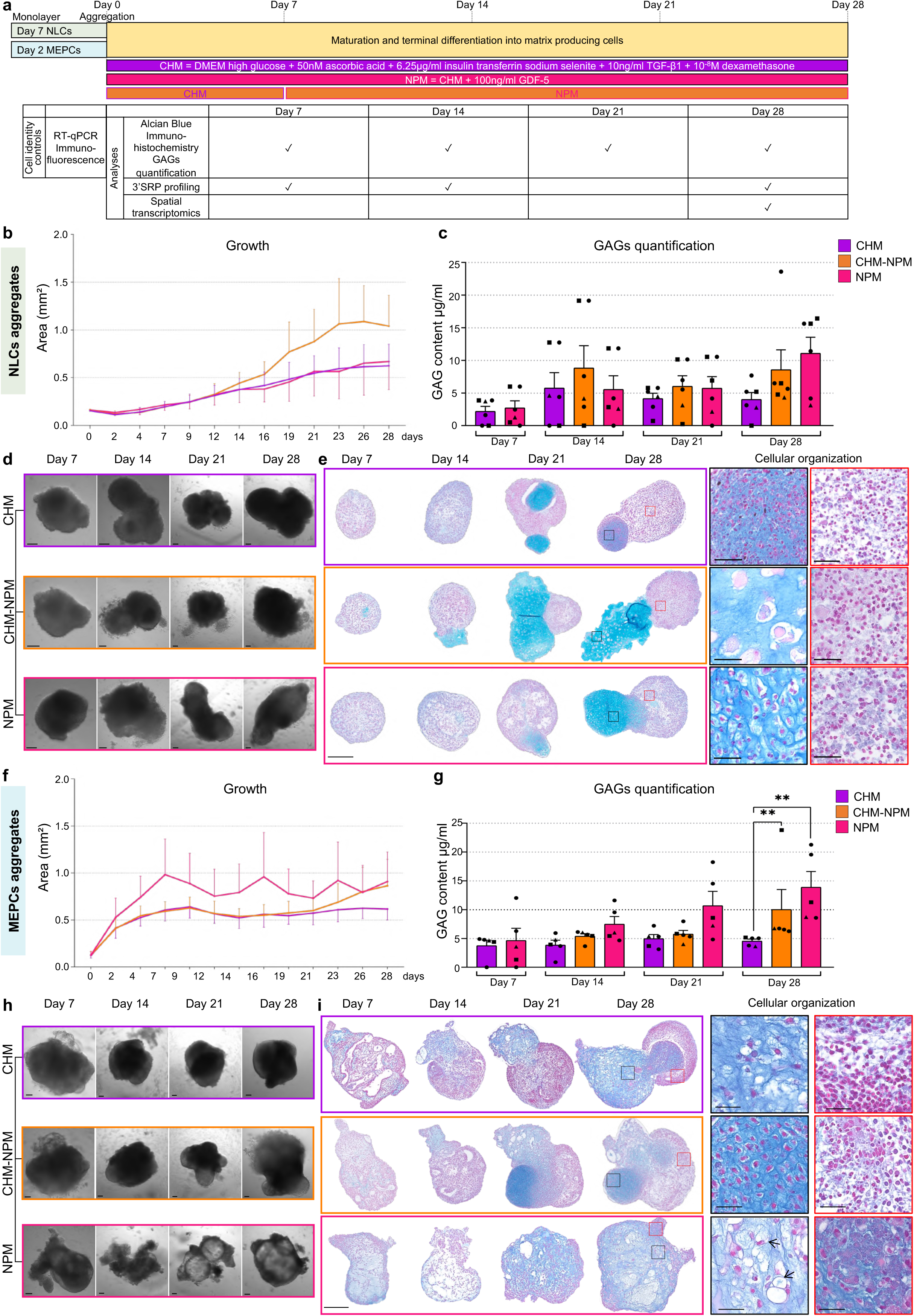
NLCs- and MEPCs-derived aggregates differentiate into matrix-producing cells with distinct structural organization in long-term 3D culture containing TGF-β1 and GDF-5. (a) Schematic of the aggregation-based differentiation protocol. The CHM condition, containing ascorbic acid, dexamethasone and TGF-β1 was compared to the NPM condition, consisting of CHM supplemented with GDF-5, and to the CHM-NPM condition involving an initial chondrogenic phase (7 days) followed by prolonged culture in NPM (21 days). (b-i) Characterization of aggregate differentiation. (b, f) Aggregate growth over time. Mean area ± s.e.m. is shown for ≥14 aggregates from five biological replicates (NLCs, three hiPSC lines) and six biological replicates (MEPCs, three hiPSC lines). (c, g) GAGs content quantification. Data represent mean ± s.e.m. from five independent experiments for NLCs and six for MEPCs; each experiment was a pool of four aggregates. 3 distinct hiPSC lines are indicated by individual symbols. Statistical significance was assessed using the Mann-Whitney test (**p < 0.01). (d, h) Brightfield images of the aggregates. Scale bar = 100µm. (e, i) Alcian blue staining on day 7, 14, 21 and 28 in each culture condition. Nuclei and GAGs are stained in blue-purple and blue, respectively. Scale bar = 250μm. Insets are showing higher magnification of cell-rich structures (red square) and ECM-rich structures (black square), and arrows point to vacuole-like structures in MEPCs_NPM aggregate. Scale bars in the insets = 50μm.

### Nucleopulpogenic medium promotes early notochordal gene regulatory networks activation in aggregates derived from mesendoderm progenitors

To examine early transcriptional states acquired in 3D culture, bulk RNA sequencing was performed at day 7 on aggregates generated either from MEPCs or from NLCs, cultured under CHM or NPM conditions. The resulting gene expression profiles were compared with a curated set of NLCs-specific genes, defined as co-regulated markers in monolayer differentiation^11^. Unsupervised heatmap analysis revealed 3 distinct expression patterns, reflecting both the cellular origin of the aggregates and the influence of culture conditions (Fig. 4a). Pattern 1 was characterized by elevated expression of core NLCs markers, including *NOGGIN, HOPX, TBXT* and *SHH*. This signature was most prominent in NLCs-derived aggregates, consistent with the persistence of their initial transcriptional identity following aggregation. These markers were also detectable in MEPCs-derived aggregates cultured under both CHM and NPM conditions, indicating early acquisition of a notochordal program. Pattern 2 was specific to MEPCs-derived aggregates, regardless of culture condition. The signature was enriched for the *SOX9* transcriptional regulator involved in matrix production as well as ECM-associated genes, including *COL2A1* and *COL9A1*^19,22^. Remarkably, the most extensive pattern 3, was uniquely enriched in MEPCs_NPM aggregates. This signature included previously described notochordal-specific targets of *SOX9* (*SERPINE2*, *ITM2A, PRELP* and *NEDD9*), and the *CREB3L2* regulon associated with NLCs differentiation^11^. In addition, MEPCs_NPM aggregates selectively expressed the surface markers *CD109 and ALCAM*, both associated with NLCs and human fetal notochordal cells^11,64^ (Fig. 4a). In line with this pattern, expression of notochord-associated markers such as *KRT8*, *KRT18*, *SOX6*, as well as *NEDD9, PRELP,* and *ITM2A,* was significantly elevated in MEPCs_NPM aggregates (Fig. 4b). To evaluate lineage specificity, we examined sclerotome and neural crest lineage specific marker expression. MEPCs_NPM aggregates showed minimal expression at day 7, confirming preferential commitment toward a notochordal fate over alternative mesodermal or ectodermal lineages, whereas CHM conditions showed higher levels of these later lineage markers (Fig. 4b). Together, these data indicate that MEPCs_NPM aggregates acquire an early notochord-like transcriptional state by day 7, marked by activation of NLCs-associated gene regulatory programs.

**Figure 4.**
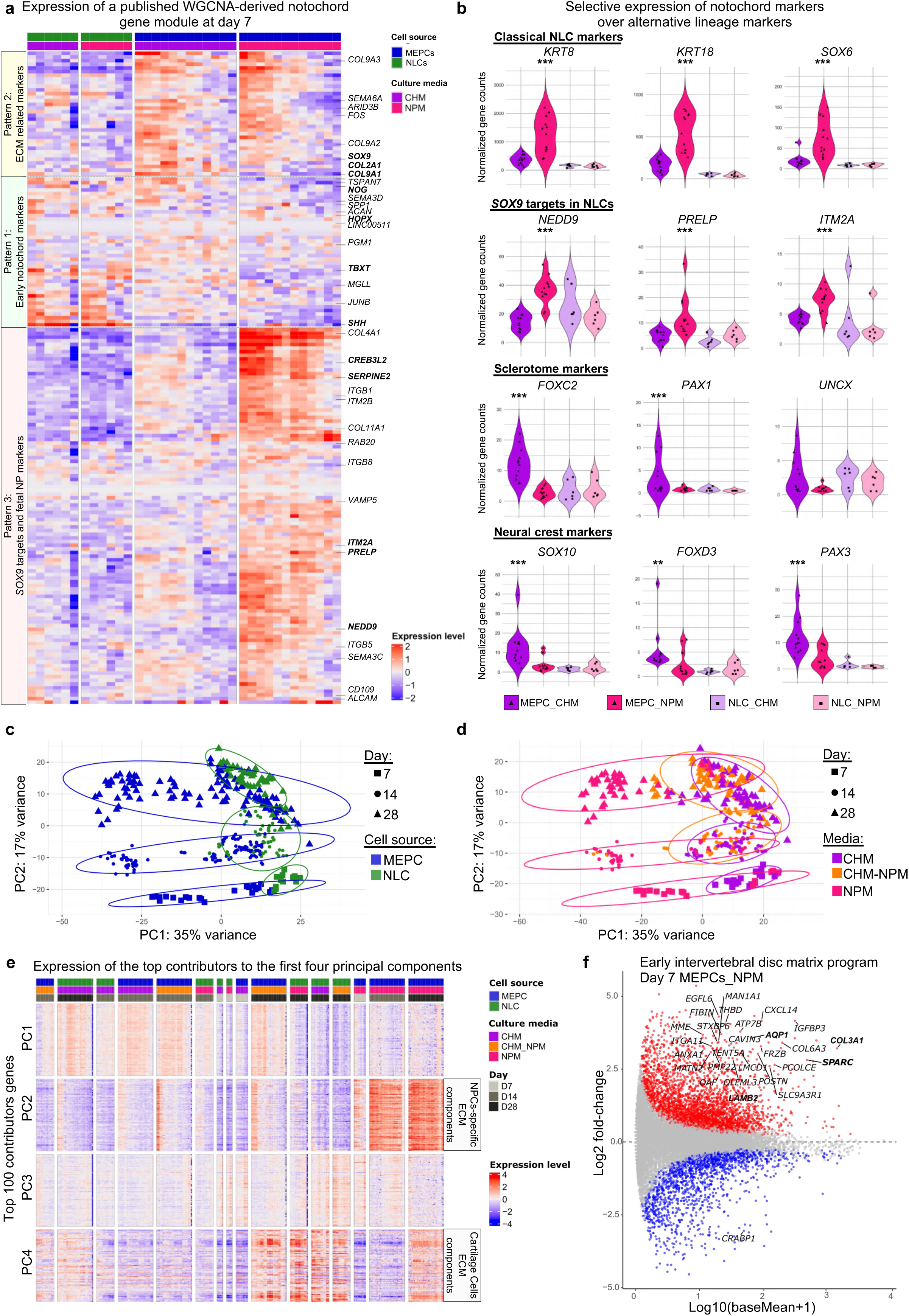
MEPCs_NPM aggregates acquire a specific NLCs-like transcriptomic profile at day 7. (a) Heatmap showing the expression of ECM-related genes, NLC markers, and *SOX9* targets and juvenile NP markers in aggregates at day 7. The marker list was derived from the WGCNA performed in Warin al., 2024.^11^ (b) Violin plots showing normalized gene counts from bulk RNA-seq for classical NLC markers (*KRT8*, *KRT18*, *SOX6*), SOX9 target genes (*NEDD9*, *PRELP*, *ITM2A*), sclerotome (*FOXC2*, *PAX1*, *UNCX*) and neural crest markers (*SOX10*, *FOXD3*, *PAX3*). Each dot represents an individual sample, shape-coded by cell source. Statistical significance was assessed using the Wald test, and p-values were adjusted for multiple testing using the Benjamini-Hochberg procedure. (**p < 0.001; ***p < 0.0001). (c-d) PCA of bulk RNA-seq data from MEPCs- and NLCs-derived aggregates over the course of differentiation. Each point represents one sample (one aggregate replicate), positioned by the first two principal components. Samples are color-coded by cell source (c) or culture medium (d); symbol shape indicates time point. (e) Heatmap of the selected top 100 genes (with the highest absolute loadings) contributing to PC1-PC4. (f) MAplot showing differential gene expression between day 7 MEPCs_NPM aggregates and all other samples collected at day 7. Each point represents a gene, plotted according to its mean expression level across all D7 samples (average log expression; x-axis) and its log2 fold change in MEPCs_NPM relative to other D7 conditions (y-axis). Genes with a mean expression value greater than 1 on the x-axis are highlighted, emphasizing transcriptional changes among robustly expressed genes. Significantly upregulated and downregulated genes (adjusted p ≤ 0.05) are indicated in color, while non-significant genes are shown in grey. In the heatmaps, each row represents a gene and each column a sample; expression levels are color-coded from low (blue) to high (red).

### Long-term 3D culture sustained by TGF-β1 and GDF-5 promotes differentiation of mesendoderm progenitors into NP-like cells

Based on ECM deposition and morphological features, divergence in differentiation trajectories across conditions was evident by day 14, while NPC-like features became apparent in MEPCs_NPM aggregates by day 28. Thus, to capture temporal transcriptional changes during differentiation, bulk profiling was further performed across all culture conditions on days 14 and 28. Principal component analysis (PCA) revealed that samples segregated by cell source along PC1 (35% of variance) with PC2 (17% of variance) reflecting time-dependent variation in both MEPCs- and NLCs-derived aggregates (Fig. 4c). A subset of MEPCs-derived aggregates showed partial overlap (Fig. 4c). When stratified by culture condition, the overlap in PCA projection was largely attributable to CHM and CHM-NPM samples (Fig. 4d). Remarkably, MEPCs_NPM samples occupied distinct regions of the PCA space at each time point while NLCs-derived samples displayed comparatively limited dispersion across conditions.

Because PCA provides a low-dimensional projection of high-dimensional transcriptomic data, interpretation was further guided by examining the genes contributing most strongly to variance along PC1 to PC4 as well as their expression patterns across samples (Fig. 4e; Table S2). Heatmap analysis revealed that PC2 captured a transcriptional program selectively enriched in MEPCs_NPM aggregates across days 7, 14, and 28. Genes contributing to PC2 were predominantly associated with ECM components, including *COL5A1* and *COL6A2,* which are known to be expressed in native NPCs derived from rabbit and mouse discs and are commonly notochordal cell-rich in a healthy state^65,66^. In contrast, PC4 reflected a shared transcriptomic program across all aggregates by day 28, regardless of initial cell source or culture condition, enriched in general chondrogenic ECM genes such as *COL2A1* and *ACAN,* which are commonly expressed by both articular chondrocytes and NPCs. To evaluate IVD compartment specificity, we used gene sets for NP and annulus fibrosus (AF) tissues (see Materials and Methods, Gitlink). This comparison revealed specific enrichment of NP-associated genes in MEPCs_NPM samples, whereas AF-associated markers remained minimal across all conditions and time points (Fig. S3a). Of the NP-associated genes, 30 were among the top 100 contributors to PC2 and were selectively expressed in MEPCs_NPM aggregates at days 14 and 28, with partial expression detectable as early as day 7 for *SPARC* gene, which interacts with the ECM, regulates cell-matrix interactions, and supports disc integrity, and *Collagen VI* that is enriched in fetus and young NP^67,68^(Fig. 4f, Fig. S3b). At this early time point, MEPCs_NPM aggregates were enriched for GO terms related to ECM organization, connective tissue, and cartilage development, indicating early activation of matrix-associated biological programs (Fig. S4a-c). By comparison, MEPCs_CHM conditions enhanced a neural-associated transcriptional program as revealed by differential expression and GO analyses (Fig. S5a-d), consistent with previous reports^69^. A similar but weaker trend was observed under CHM-NPM conditions. Together, these analyses indicate that MEPCs_NPM aggregates activate a distinct NP- and matrix-associated transcriptional program, while a general ECM signature emerges across all conditions at later stages. Subsequent analyses, therefore, focused on NPM-cultured aggregates.

### NP-specific gene regulatory networks and matrisome profile define structural maturation of MEPCs_NPM aggregates

We investigated the molecular networks across all media conditions and cell sources to characterize the emergence of NP-specific features in aggregates at day 28. WGCNA analysis identified five major modules of co-expressed genes (Fig. S5e, [Gitlink](https://github.com/w-julie/3D-cell-paper/tree/main/bulkRNAseq/5-WGCNA)). Interestingly, a distinct module comprising 596 genes was selectively enriched in MEPCs_NPM aggregates. Comparison analysis with the NP-associated gene list, revealed 104 shared genes, validating the biological relevance of this module to NP identity (Fig. 5a). Functional annotation of this NP-enriched module revealed a strong representation of ECM-related genes upregulated in MEPCs_NPM aggregates from day 7 to day 28 (Fig. 5b). Remarkably, numerous components of NP core matrisome^15,70,71^, including collagens (*COL1A1*, *COL5A1*, *COL6A2*, *COL4A1*, *COL4A2*) and glycoproteins (*MFGE8*, *CRISPLD2*, *SLIT3*, *PCOLCE*, *FBLN1*), were prominently represented. Matrisome-associated genes were also identified, encompassing ECM-affiliated proteins (*SEMA7A*, *PLXNB2*, *PLXND1*, *ANXA6*, *PLXNA3*), ECM regulators (*SERPINE2*, *CST3*, *ADAM33*, *ADAMTS2*, *BMP1*), and secreted factors implicated in ECM organization and signalling (*ANGPTL7*, *TGFB1*, *FSTL3*, *EGFL7*, *FST*). In comparison, these ECM-related genes showed systematically low expression in NLCs-derived aggregates (Fig. S6). Consistent with these network-level findings, GO enrichment demonstrated that the transcriptome of MEPCs_NPM aggregates at day 28 was significantly enriched for biological processes related to ECM organization (GO:0030198), collagen metabolic processes (GO:0032963), connective tissue development (GO:0061448), and cartilage development (GO:0051216) (Fig. S4c, [Gitlink](https://github.com/w-julie/3D-cell-paper/blob/main/bulkRNAseq/5-WGCNA/WGCNA_18_moduleTurquoise_enrichGO.csv)). Together, these analyses demonstrate that, from day 7, MEPCs_NPM aggregates adopt a coordinated ECM program characteristic of NP tissue architecture and the IVD ECM landscape essential for homeostasis.

**Figure 5.**
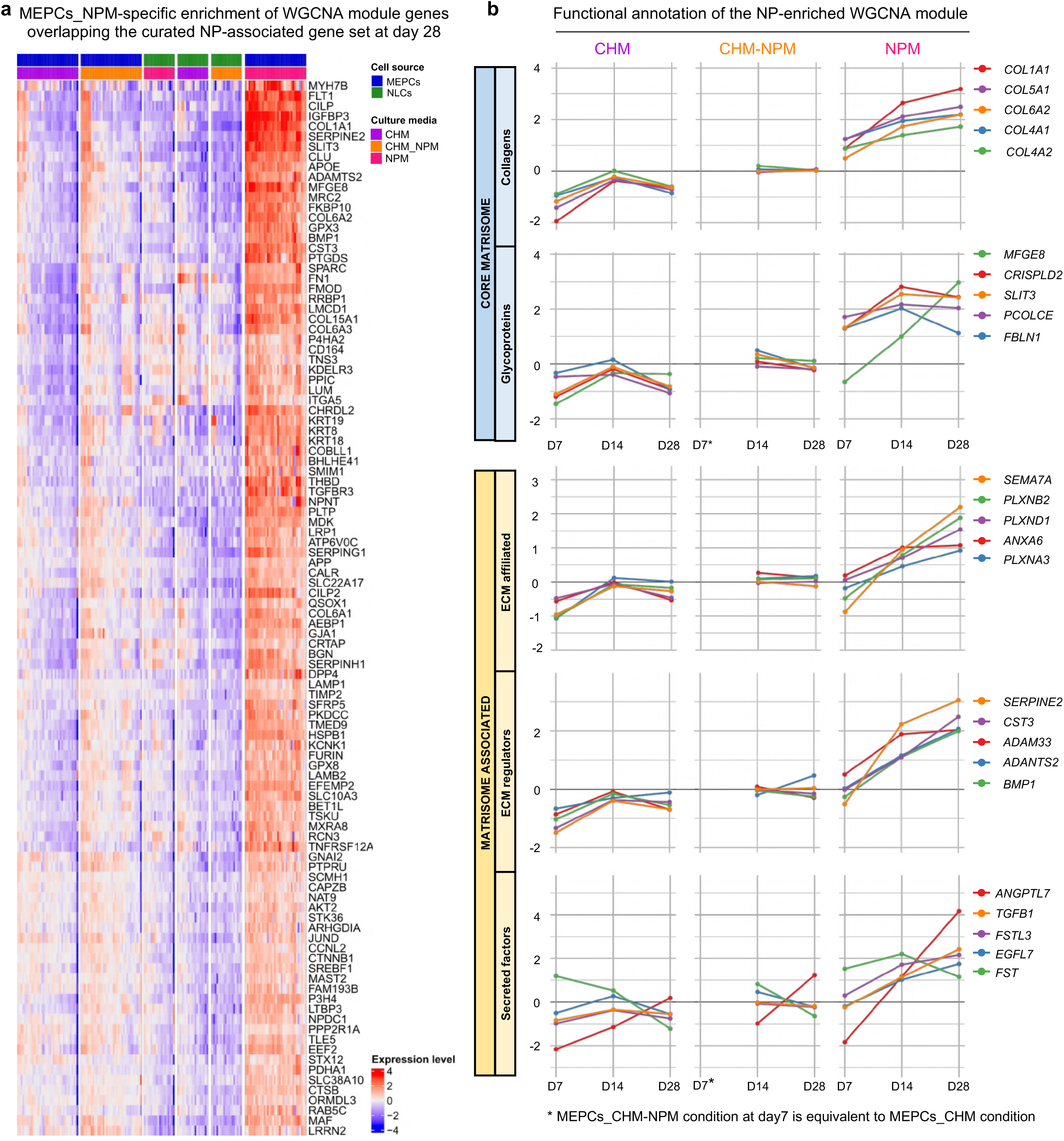
MEPCs_NPM aggregates acquire a transcriptomic profile similar to native NP cells. (a) Heatmap showing the expression of NP markers from the WGCNA module associated with MEPCs_NPM aggregates at day 28. Each row represents a gene and each column a sample; expression levels are color-coded from low (blue) to high (red). (b) Temporal expression profiles of key genes from the WGCNA-identified module across differentiation (days 7, 14, and 28). Line plots display centered expression dynamics in MEPCs_CHM aggregates (left), CHM-NPM (middle), and NPM (right). Genes are grouped by functional categories: core matrisome (collagens, glycoproteins), ECM-affiliated proteins, matrisome-associated proteins, and growth factor-related factors.

To validate NP-associated transcriptional signature at the protein level, we examined the expression and spatial distribution of notochordal and NP cell markers in aggregates across all conditions and cell sources. We focused on markers shared between these two lineages, including SOX9, KRT8, CD109 and CD166 (ALCAM), as well as the angiopoietin receptor TIE2 (TEK) expressed in disc progenitor populations^11,15,64,71,72^. Consistent with their broad lineage expression^73,74^, *SOX9-* and *CD166*-positive cells were detected across conditions (Figs. 6, S7 and data not shown). Their presence in NLCs-derived aggregates with neural-associated transcriptional program confirms their limited NP-specificity, necessitating combinatorial marker analysis. In NLCs-derived aggregates KRT8, CD109 and TIE2 immunostaining remained low, sparse and spatially confined (Fig. S7a,b), consistent with weak transcript-level expression. By contrast, MEPCs_NPM aggregates exhibiting high transcript-level expression showed widespread markers immunostaining first detected at day 14 and increasing in intensity and distribution by day 28 (Fig. 6a,b), underscoring progressive acquisition of an NP-like phenotype.

**Figure 6.**
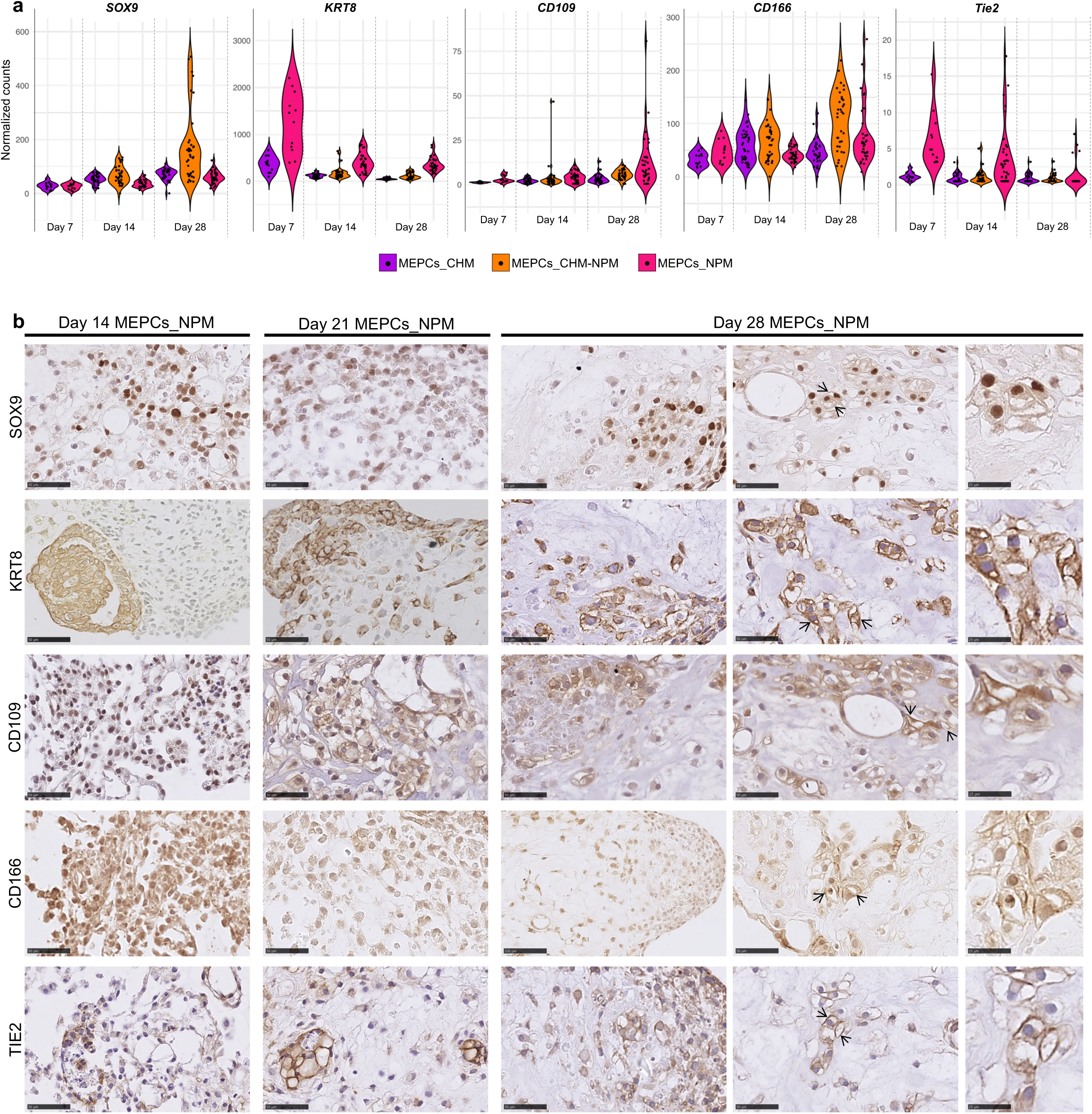
Transcriptional and protein-level validation of NLC/NP markers in MEPCs_NPM aggregates. (a) Violin plots showing normalized gene counts from bulk RNA-seq for NLC/NP markers (*SOX9*, *KRT8*, *CD109*, *CD166, Tie2*). Each dot represents an individual sample, shape-coded by cell source. (b) Representative immunohistochemistry of NP markers at day 14, 21, and 28 in MEPCs_NPM aggregates. At day 28, two distinct area of the aggregate are showing morphology suggestive of vacuole-like structures (arrows) with insets showing higher magnification. Scale bars, 50 µm; insets, 25 µm. Images are representative of at least 3 independent experiments and samples.

CD109 immunostaining corroborated the maturation process in MEPCs_NPM aggregates, revealing vacuole-like morphological features also detected by Alcian blue staining at day 28 (Figs. 3h, 6b). To investigate the molecular basis of vacuolation in 3D culture, we used gene sets associated with lysosome-related organelles (LROs) (see Materials and Methods, Gitlink) for comparative analysis with differentially expressed genes, which demonstrated a specific and statistically significant enrichment of LRO-associated genes in MEPCs_NPM aggregates at day 28 (Fig. 7a). Consistent with the formation of specialized intracellular compartments, and vacuole biology, upregulated genes included regulators of vesicle trafficking and membrane dynamics (*DNM1* and *DNM2*^75^, *RAB32*^76^, *RAB5C*^71^ and *VAMP8*) and genes linked to organelle homeostasis and ion regulation (*NBEAL2*, *OCA2* and *OSTM1*). Notably, *LAMP1*, a canonical lysosomal membrane protein, previously shown to localize at the vacuole membrane of notochordal cells in the mouse embryo^8,71^ was strongly enriched. Confirming the high expression at the transcript level, immunostaining for LAMP1 and CAV1^18,77^ were detected in vacuolar structures specifically in MEPCs_NPM aggregates at day 28 (Fig. 7b). In contrast, NLCs aggregates were composed of smaller, densely packed cells lacking vacuolated morphology (data not shown). Overall, these results link the characteristic vacuolar morphology in MEPCs_NPM aggregates to the selective activation of an LRO-associated transcriptional program, consistent with maturation-associated characteristics of notochordal progenitors and NP cells.

**Figure 7.**
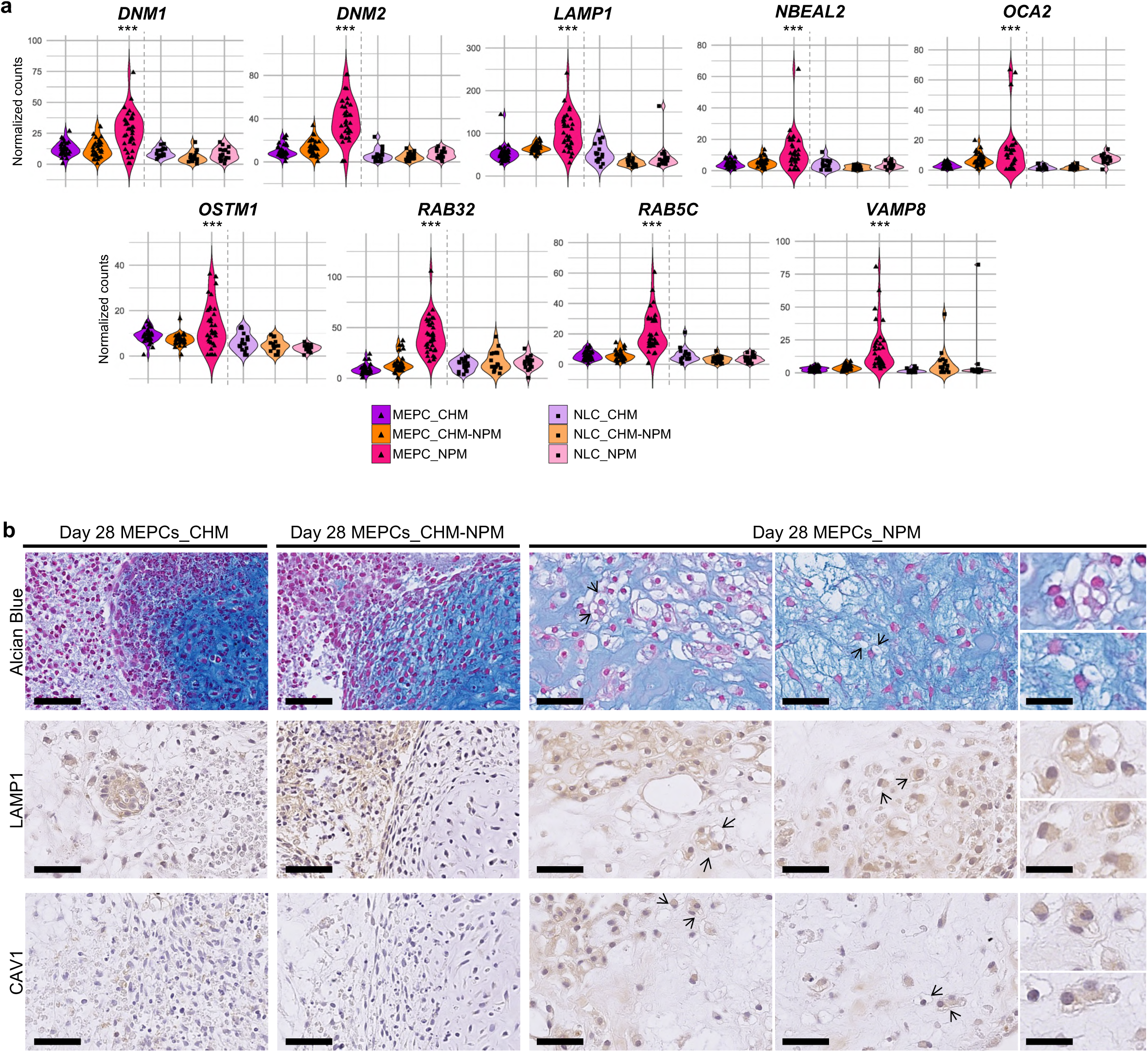
MEPCs_NPM aggregates differentially express vacuole-related markers at day 28. (a) Violin plots showing normalized gene counts from bulk RNA-seq at day 28 for vacuole-related makers (i.e. gene regulators of vesicle trafficking and membrane dynamics). *DNM1* and *DNM2* is involved in the control of vesicle formation during endocytosis and *RAB32*, *RAB5C* and VAMP8 membrane in vesicle fusion. Each dot represents an individual sample, shape-coded by cell source. Statistical significance was assessed using the Mann–Whitney test (***p < 0.0001). (b) Representative Alcian blue staining and immunohistochemistry for markers LAMP1 and CAV1 at day 28 in MEPCs aggregates under each culture condition. In the NPM condition, two distinct area of the aggregate is shown, revealing extensive vacuolar morphological features (arrows) with insets showing higher magnification. Scale bars, 50 µm; insets, 25 µm. Images are representative of at least 3 independent experiments and sample.

### Spatial gene expression mapping identifies dual notochordal and nucleus pulposus programs in MEPCs_NPM aggregates

To provide a functional overview of the transcriptional differences between conditions, we performed gene set variation analysis (GSVA, [Gitlink](https://github.com/w-julie/3D-cell-paper/tree/main/bulkRNAseq/7-GSVA)). MEPCs_NPM aggregates exhibited a coordinated upregulation of cartilage- and ECM-associated programs, encompassing matrix organization, collagen and proteoglycan metabolism, and cartilage-associated developmental processes (Fig. 8a-e). This transcriptional program was consistently observed across biological replicates, indicating a robust and progressive acquisition of an ECM- and cartilage-associated phenotype in MEPCs_NPM aggregates over time. Finally, we performed spatial transcriptomic profiling of MEPCs_NPM aggregates and applied gene set-based scoring using NC, NP and LRO signatures to analyze their spatial organization at day 28 (Fig. 8f, g). NP- and NC-associated transcriptional programs were enriched across the aggregates and displayed extensive spatial overlap, while the LRO-associated genes were detected in approximately 20% of scored cells (Fig. 8h). Spatially resolved transcriptomic profiling revealed the expression of key NP-associated genes with characterized functional roles, including *SPARC* and *SERPINE2 (*Fig. 8i), whose loss or reduced expression has been linked to premature intervertebral disc degeneration in mouse and human, respectively^67,78^. *GLUT1*, a marker of human NP-associated marker of hypoxic adaptation shown upregulated during disc degeneration^14^, was detected in a subset of cells (Fig. 8i), further supporting the relevance of this model to study the transition of NCs to a mature phenotype, as well as NP formation and homeostasis.

**Figure 8.**
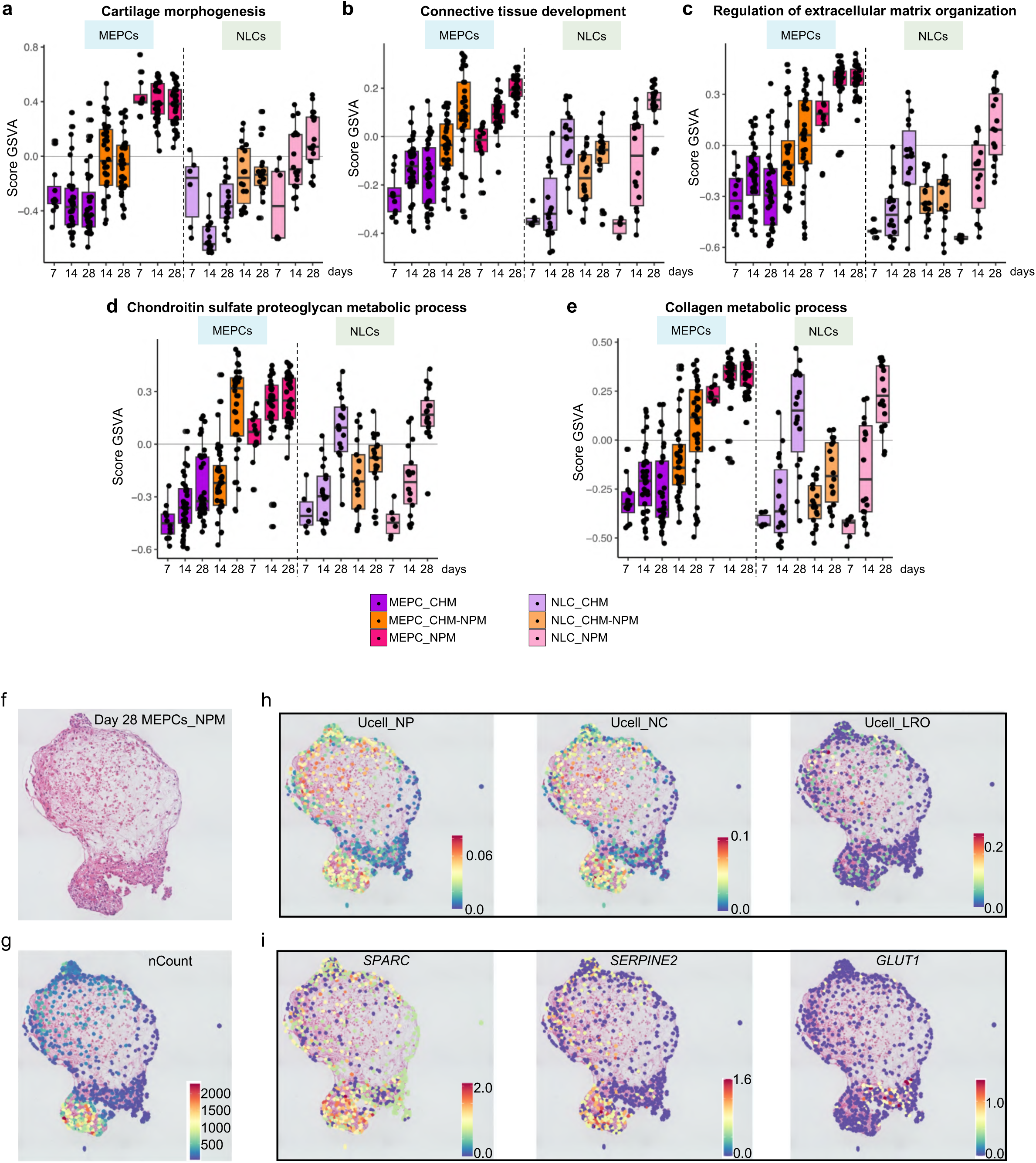
Spatial and temporal characterization of notochordal and NP-associated programs in MEPCs_NPM aggregates. (a-e) Progressive enrichment of cartilage- and extracellular matrix-associated biological processes in MEPCs-derived aggregates in NPM. GSVA scores for cartilage morphogenesis (a), connective tissue development (b), regulation of extracellular matrix organization (c), chondroitin sulfate proteoglycan metabolic process (d), and collagen metabolic process (e) across the different cell source, culture condition and time points. MEPCs-derived aggregates in NPM condition showed a progressive increase in pathway activity over time, with consistent distributions across biological replicates. (f) Representative histological image of MEPCs_NPM aggregate at day 28 and (g) corresponding spatial transcriptomic section used for spatial gene expression profiling. (h) Spatial gene set-based UCell scoring of curated NP, NC, and LRO transcriptional signatures. (i) Spatial expression patterns of selected NP-associated genes (*SPARC*, *SERPINE2*; *GLUT1*). LRO: lysosome-related organelles; NC: notochordal cell; NP: nucleus-pulposus;

## Discussion

In this study, we demonstrate that hiPSCs-derived MEPCs share transcriptional similarity with early embryonic organizer and axial mesendoderm-related populations and can be differentiated through a developmentally relevant trajectory into NP-like cells. 3D self-organization induces endogenous *NOTO* expression and enhances MEPCs commitment to early notochordal fate. Using multi-omic and morphological concordance to canonical NC and NP-like criteria, we demonstrated that long-term aggregate culture with TGF-β1 and GDF-5 promotes the progressive maturation of MEPCs into NLCs and into functional NP-like cells which exhibit robust matrix secretory phenotype and typical vacuolar morphology, therefore recapitulating the notochord-to-NP developmental continuum *in vitro*. This 3D culture system induces gene regulatory programs characteristic of in *vivo* differentiation. Spatial analysis reveals co-existing notochordal and NP programs and localized LROs signatures in day 28 aggregates, supporting further phenotypic maturation and reflecting morphological and histological features of the native healthy NP niche.

During vertebrate embryogenesis, the notochord originates from the organizer regions, which transiently harbour bipotent mesendodermal populations at early gastrulation stages. Organizer regions integrate multiple signaling pathways to coordinate axial patterning and lineage allocation, yet these processes remain largely inaccessible to functional experimentation^1^. Recent scRNAseq transcriptomic atlases in mouse embryos have provided unprecedented molecular resolution of these organizer-associated cells around embryonic day E7-E7.5, yet the scarcity of these cells, with fewer than 50 typically captured across multiple embryos^53^, poses a major challenge for detailed functional characterization. In humans, studying such rare populations therefore requires innovative strategies to faithfully recapitulate their properties *in vitro*. Here, we identify MEPCs, as a transcriptionally defined progenitors that align with organizer and axial mesendoderm-related signature *in vivo*^10,12,58,79^. The SCENIC-inferred regulons *GSC*, *KLF8*, *LHX1*, *MIXL1*, all implicated in organizer and axial mesendoderm function *in vivo*^53–55^, define a conserved regulatory network that is absent from other progenitor cell clusters, including PS and NMP, further underscoring the distinct molecular identity of MEPCs. Furthermore, predicted interactions between *LHX1*, *OTX2*, *FOXA2*, *GSC and HHEX* factors could provide, upon experimental validation, new insights into the regulatory mechanisms governing early lineage specification during human development^1,10,58^. These cells constitute a relevant and tractable model for investigating cell behaviour and function, enabling mechanistic exploration of early developmental programs and bridging a critical gap where direct experimentation is not feasible.

In the present study, this organizer-like state is deliberately used as an entry point to access the notochordal lineage and to examine the subsequent transition toward NP-like identities. Starting from this defined progenitor state, our study provides the first evidence that notochordal identity can be induced endogenously through 3D self-aggregation of MEPCs, without requiring ectopic transcription factor expression, unlike most existing *in vitro* approaches that rely on forced expression of *NOTO, BRACHYURY or FOXA2* to generate NLCs^11,36,37^, also reflecting the limits of activating the notochordal regulatory program under conventional two-dimensional culture conditions^39,42^. A recent study demonstrated that these MEPCs, selected as a cell source and co-injected with an instructive hydrogel in degenerated discs in an experimental dog model, engrafted and adopted a functional matrix-producing phenotype expressing *ACAN* and *COL2*^80^. Nevertheless, *in vitro* maturation of NLCs toward NP-like cells has remained a complex issue. A major bottleneck is the inability to expand native notochordal cells in a stable manner^81^, which restricts direct investigation of the maturation transcriptional program. Leveraging prior evidence linking TGF-β and GDF5 signaling to IVD development^82,83^ we show that MEPCs aggregation in combination with these growth factors creates a permissive developmental microenvironment. This was marked by formation of vacuole-like structures, a defining feature of notochord maturation rarely recapitulated in human *in vitro* model^47^, and upregulation of NP-associated markers. In this study combinatorial markers (e.g. SOX9, KRT8, CD166, CD109) previously associated with native NP cells improved the specificity of NP-like cells’ identification. Time-resolved, large-scale transcriptomic profiling revealed progressive activation of NP gene regulatory networks alongside coordinated ECM and phenotypic maturation. Multi-timepoint sampling across biological replicates established model robustness and resolved dynamic transcriptional states. The presence of LROs signatures and vacuole-associated markers provides further evidence that this model captures a developmental continuum encompassing immature NLCs, vacuolated notochordal states, and NP-like cells, which could be further dissected using time-resolved single-cell omics. Altogether our results demonstrate maturation and NP-like physiology with functional attributes inferred from increasing NP hallmarks during differentiation, NP-associated transcriptomic signatures and matrisome modules, robust GAG deposition and vacuole-like morphology. Direct functional assays, such as measurements of compressive mechanics, swelling pressure, or osmotically driven responses, will confirm NP-specific physiological and biomechanical performance of the aggregates.

TGF-β signaling is a central regulator of notochordal cell maturation and NP function^26^. However, excessive or dysregulated TGF-β activity has also been implicated in IVD degeneration, highlighting the need for tight spatial and temporal control of this pathway. The detection of the TGF-β antagonist *LEFTY2* in MEPCs_NMP aggregates suggests the presence of a potential intrinsic negative feedback mechanism that may contribute to signaling homeostasis. Consistent with this hypothesis, the ECM composition of MEPCs_NPM aggregates reflects an intermediate developmental state between fetal or adult phenotype. Several matrix components preferentially expressed during human fetal NP development, including *FMOD*, *BGN*, *PCOLCE*, *COL6A1* and *COL6A3*, were robustly detected, whereas others such as *MFGE8* increased progressively from day 7 to day 28, suggesting ongoing matrix maturation^84^. At the same time, genes more highly expressed in non-degenerated adult NP, including *VCAN*, *FN1*, *LUM* and *CILP*, were also induced, indicating convergence toward an adult-like NP matrix signature^84^. Notably, several of these matrix components have been directly linked to disc integrity and protection. Biglycan deficiency in mice leads to premature IVD degeneration^85^, while versican has been shown to be enriched in the human NP relative to the AF, and to promote NP cell proliferation and adhesion *in vitro*^86^. Taken together, these findings indicate that MEPCs-derived aggregates establish a controlled ECM environment combining fetal-associated plasticity with adult NP homeostatic features, rather than progressing toward unchecked TGF-β-driven fibrotic matrix deposition often seen when disc integrity is compromised^26^. Such balanced regulation may be critical for preventing degenerative trajectories. Finally, to define the model’s transcriptomic divergence from native human NP cell states and to quantitatively established its high fidelity, direct integration of the aggregate transcriptomes with matched fetal human NP single-cell datasets across developmental stages up to juvenile stage is required. However, publicly available IVD-specific developmental transcriptomic atlases have only recently emerged, limiting benchmarking possibilities.

Beyond intrinsic cell fate specification, the coexistence of NLCs and NP-like cells within the 3D aggregates indicates that this system recapitulates key aspects of the native healthy NP niche. NCs have been proposed to exert trophic and protective effects on NP cells through paracrine signaling and matrix-mediated interactions^20,25,27–30^, yet these processes have been difficult to interrogate directly in human tissues. This robust, developmentally and physiologically relevant human *in vitro* model provides a platform to investigate such intercellular dialogue with the NP, and to determine how NCs loss contribute to disc deterioration, while evaluating the potential protective role of NCs in maintaining disc integrity. Beyond advancing our understanding of NC biology and physiology, this model is instrumental in guiding the selection of appropriate cell sources for preclinical evaluation, and facilitating the development of novel disease-modifying strategies for clinical translation.

## Methods

### Ethics statement

The hiPSC line^a^ was obtained from commercial Life Technologies. The hiPSC line^b^ (hPSCreg Name: REGUi016-D 4F_PBMC_Sv190) was originally derived from patient cells collected at Inselspital Hospital of the University of Bern, Switzerland (Swiss ethics approval no. 2019-00097). The third hiPSC line corresponds to hPSCreg Name: LIMFRi001-A and the fourth was developed in the iPSC core facility Nantes.

### 1. Human induced pluripotent stem cell culture

Stem cell culture was achieved in line with the ISSCR guidelines^1^. HiPSC lines used in this study were maintained on ESC-qualified Matrigel (#354277, Corning), coated plates with mTeSR1 media (#85850, Stem cell technology). All cell lines were cultured at 37°C, under normoxic conditions (20% O2, 5% CO2) and culture media was daily replaced. HiPSC were passaged every 4 days at a 1:6 split ratio using passaging solution XF (#130-104-688, Miltenyi) for 2 min, at room temperature for dissociation into cell clusters. Cells were grown for 10 passages maximum. A new vial of the cell bank was used when cell reach P30. The MycoAlert kit (#LT07-318, Lonza) was employed to confirm the absence of mycoplasma in all cell lines.

### 2. Notochordal-like cells differentiation

Differentiation protocol followed two-steps as described in Warin et al. 2024^2^: (1) experiment started with the seeding of single hiPSC at 20.000 cells/cm2 on Laminin-521 (#354224, Corning)-coated plates in mTseR1 medium supplemented with rock inhibitor (Y27632, #1683, Axon MedChem) for 24 hours; (2) cells were then cultured in specific media for mesendoderm progenitor specification [DMEM/F12 (#11330, Life Technologies) supplemented with 1% N2 (#17502048, Gibco), 1% B27 (#17504044, Gibco), 1% non-essential amino acids (#11140, Life Technologies), 1% GlutaMAX (#35050, Gibco), 0.001 mM 2-mercaptoethanol (#31350, Sigma), 50 mg/ml bovine serum albumin (#A-1595, Sigma) and 3µM CHIR99021 (#1386, Axon Medchem)]. The mesendoderm progenitors are then transfected with 1,5µg of synthetic NOTO mRNA using lipofectamine RNAimax (#13778075, Life Technologies). The transfection is either performed in monolayer, or for 4 hours in suspension for subsequent 3D culture. After 4 hours of transfection, 35 000 cells (or 70 000 cells for the transfected condition) were aggregated in N2B27 + 3µM CHIR + 100ng/ml noggin (#130-103-456, Miltenyi) + 10µM Y27632 by centrifugation (50g for 5 minutes) in 96-well plate round bottom low attachment and placed in the incubator at 37°C, and 5% CO2.

### 3. Nucleopulpogenic differentiation

After dissociation in TryplE (#12604021, Life Technologies), 35 000 MEPCs or NLCs were aggregated in N2B27 + 3µM CHIR + 100ng/ml noggin + 10µM Y27632 by centrifugation (50g for 5 minutes) in 96-well plate round bottom low attachment and placed in a hypoxia chamber (37°C, 5% O2, 5% CO2). Culture media conditions were previously described in Colombier et al. 2016^3^ and are defined as follow. The control group was cultivated in the presence of a basal medium (BM) composed of DMEM high glucose (#61968-026, Gibco) supplemented in 1% penicillin/streptomycin (#15140-122, Gibco), 50nM stable acid ascorbic (#A4034, Sigma-Aldrich), and 6.25 µg/ml Insulin-transferrin-sodium selenite (#I1884-1VL, Sigma). For the chondrogenic condition (CHM) the BM was supplemented with 10ng/ml recombinant human TGF-β1 (#100-21C, Peprotech) and 10^-8^M dexamethasone (#D4902, Sigma). For the nucleopulpogenic condition (NPM), the CHM was supplemented with 100ng/ml recombinant human GDF-5 (#120-01, Peprotech). A third condition was evaluated (CHM-NPM) involving sequential induction of CHM and NPM media with an initial chondrogenic phase (7 days) followed by prolonged culture in NPM (up to 28 days). The media were refreshed three times a week.

### 4. RNA extraction and RT-qPCR

Total RNA was extracted from pooled aggregates with the Nucleospin II RNA Kit XS (#740902.50, Macherey Nagel) and one microgram was reverse transcribed using Superscript III First Strand synthesis kit (#11752, Life technologies). Quantitative RT-qPCR experiments were performed using TaqMan or SYBR technologies with GAPDH as housekeeping gene and fold change represented using a base 2 logarithm determined by the Livak Method (Relative quantification RQ = 2^-DDCq^)^4^.

### 5. Fixation/Inclusion

Pellets were washed in PBS-Tween 0.02% three times for 5 min before 30min of fixation in 4% PFA followed by three washing steps in PBS-Tween 0.02%. The pellets were stored in PBS-Tween 0.02%-BSA 2% until further use.

For histological stainings and immunohistochemistry, aggregates were embedded in Histogel (#HG-4000-012, Epredia) before paraffin embedding and sectioning. The Histogel was heated at 90°C until it became liquid before use. The aggregates were placed in the lid of a 96-well plate (used as a mould) and all the liquid was aspirated. 80µl of liquid Histogel were then added on top of the pellet, and using a tip the aggregate was replaced at the center of the droplet and kept at RT until the Histogel solidified. The samples were then dehydrated before paraffin embedding following a classical protocol.

| Reagent | Time |
| --- | --- |
| Alcohol 80° | 1h30 |
| Alcohol 80° | 1h30 |
| Alcohol 95° | 1h30 |
| Alcohol 95° | 1h30 |
| Alcohol 100° | 1h30 |
| Alcohol 100° | 1h30 |
| Alcohol 100° | 1h30 |
| Methylcyclohexane | 1h30 |
| Methylcyclohexane | 1h30 |
| Methylcyclohexane | 1h30 |
| Paraffin | 2h15 |
| Paraffin | 2h15 |

### 6. Immunofluorescence staining of aggregates

Aggregates were permeabilized and blocked by incubation in DPBS containing 0.5% Triton X-100 and 3% bovine serum albumin (BSA) for 2 hours at room temperature under gentle agitation, with a buffer change after 1 hour. Samples were then incubated with primary antibodies diluted in DPBS supplemented with 0.5% Triton X-100 and 3% BSA for 24 hours at 4 °C under constant agitation. After incubation, samples were washed in the same buffer for 2 hours at room temperature with a buffer change halfway through. Secondary antibodies, together with 1 µM TO-PROTM-3 Iodide (#T3605, Thermo Scientific), were applied in DPBS containing 0.5% Triton X-100 and 3% BSA for 24 hours at 4 °C under agitation. From this step onward, samples were protected from light. Aggregates were then washed with DPBS containing 0.02% Tween-20 every hour for 3 hours at room temperature with agitation. Finally, samples were cleared in RapiClear 1.52 (#RC152001, Sunjin Lab) and kept at 4°C in the dark until imaging. Table S3 summarizes the antibodies information for staining.

### 7.

Histological staining and immunohistochemistry

5 μm sections were de-paraffinized, rehydrated and stained following standard protocol with Periodic Acid Schiff/Alcian Blue (PAS/AB).

For immunohistochemistry detections, after antigen retrieval, sections were permeabilized in DPBS containing 0.5% Triton X-100 for 20 min: The sections were incubated with the primary antibodies overnight at 4°C after incubation with blocking solution. The next day, the sections were incubated with secondary antibodies conjugated with biotin. Revelation was performed after 45min incubation in streptavidin-peroxidase solution and of DAB substrate (3,3’-Diaminobenzidine). Nuclei were stained with Mayer Haematoxylin, mounted with Eukitt and scanned using a Hamamatsu Nanozoomer HT (Hamamatsu Photonics KK) digital scanner at 40X magnification. Table S3 summarizes the dilution and specific buffers for retrieval for each antibody.

### 8. Single Cell RNA sequencing

Cells were dissociated and labeled using lipid-conjugated oligonucleotide barcodes (10x Genomics Multiplexing technology). A total of 40,000 cells were loaded onto the Chromium Controller (10x Genomics) for single-cell encapsulation and reverse transcription using the Chromium Single Cell 3’ Reagent Kit. Following cDNA synthesis, cleanup and amplification were performed according to the manufacturer’s protocol. Enzymatic fragmentation and adapter ligation with sample-specific indexes were conducted on double-stranded cDNA, followed by PCR amplification. Size selection was performed using a dual-sided SPRIselect magnetic bead protocol to minimize artifacts, optimize transcript coverage, and improve alignment quality. Final library concentration and fragment size distribution were assessed using Qubit (Thermo Fisher Scientific) and Bioanalyzer (Agilent). Libraries were sequenced on an Illumina NovaSeq 6000 platform with paired-end reads following the manufacturer’s recommendations.

### 9. 3’ SRP RNAseq preparation

For RNAseq analysis, the samples were stopped along the timing of the differentiation. The aggregates were stored in twice the volume of the timing (i.e day 7 in 14µl of lysis buffer) ultra pure water containing RNAse inhibitor (2,86U/µL Ambion RNase Inhibitor, Invitrogen -AM2684). Direct cell lysis of individual aggregates was performed by 3 cycles of 30min of thawing at RT, freezing at -80°C and pipetting mix, before storing.

The libraries were prepared as described in Charpentier et al.^5^ from 4µl of lysate. Briefly, the samples are tagged with universal adapters (E5V6NEXT Adaptater), well-specific barcodes and unique molecular identifiers (UMIs) (E3V6NEXT) during template-switching reverse transcription using Maxima H Minus Reverse Transcriptase. All cDNAs obtained after template-switching reaction are pooled together, purified and concentrated with single DNA & Concentrator-5 column (#ZD4013, Ozyme). Pooled and concentrated cDNAs are treated with Exonuclease I (#M0293S/L, New England Biolabs) and then amplified by single primer PCR using Advantage® 2 PCR Kit (#639206, Ozyme) and SINGV6 primer (10μM, Integrated DNA Technologies). Amplification is performed by PCR. PCR products are purified with Agencourt AMPure XP magnetics beads [0.6x] (#A63881, Beckman Coulter) and quantified specifically for dbDNA using Quantus (QuantiFluor ONE dsDNA, E4870). Amplified barcoded cDNAs are tagmented using a bead-linked transposome (Illumina). 200ng of full-length cDNAs was used as input to the Nextera™ DNA Flex Library Prep kit (#20018704, Illumina) and Nextera™ DNA CD Indexes (24 Indexes, 24 Samples) (#20018707, Illumina) according to the manufacturer’s protocol (Nextera DNA Flex Library Document, #1000000025416 v04, Illumina) with exception that the i5 primer was replaced by P5NEXTPT5 primer (5μM, Integrated DNA Technologies), the number of PCR cycles was defined at 8, and final elution of purified library was realized in 15µl RSB

(Resuspension Buffer - included Nextera™ DNA Flex Library Prep kit, Illumina). Libraries were sequenced on an Illumina NovaSeq 6000 platform

E3V6NEXT = 5’-/5Biosg/ACACTCTTTCCCTACACGACGCTCTTCCGATCT[BC6]N10T30VN-3’

5Biosg = 5’biotin

[BC6] = 6bp barcode specific to each aggregate/well

N10 = unique molecular identifier

E5V6NEXT = 5’-/5Me-isodC//iisodG//iMe-isodC/ACACTCTTTCCCTACACGACGrGrGrG-3’

SINGV6 = 5’-/5Biosg/ACACTCTTTCCCTACACGACG*C-3’

P5NEXTPT5 =

5’-AATGATACGGCGACCACCGAGATCTACACTCTTTCCCTACACGACGCTCTTCC*G*A*T*C*T*-3’

where * = phosphorothionate bonds

### 10. Bioinformatic analysis for single-cell RNAseq

Raw sequencing data were processed using Cell Ranger (v6.0.2, 10x Genomics), and FASTQ files were aligned to the human reference genome (GRCh38-2020-A). Sample demultiplexing and quality filtering were performed using Seurat (v4) ^6^. Dataset-specific thresholds for the number of detected genes and the percentage of mitochondrial gene expression were applied to exclude doublets, dead cells, and empty droplets. Data were normalized using the SCTransform method. Each dataset, including previously published data, was analyzed independently and subsequently integrated using the ComBat-seq algorithm to ensure results are robust to integration choice. The integration strategy was validated by (i) assessing dataset dispersion across hiPSC lines, (ii) comparing observed and expected cell numbers per cluster, and (iii) confirming that early timepoint samples did not cluster with later timepoints.

Dimensionality reduction was performed by PCA on highly variable genes. Unsupervised clustering was conducted using a shared nearest neighbor (SNN) graph constructed from the top principal components, based on the k-nearest neighbors (KNN) algorithm. Clusters were visualized using Uniform Manifold Approximation and Projection (UMAP). Differentially expressed genes (DEGs) were defined within each cluster as genes detected in at least 10% of the cells, with a positive average log fold-change > 0.25. Cluster identities were annotated manually based on DEGs signatures and known marker genes from the literature (Table S1). DEGs lists from each dataset were compared before and after integration using VennDiagram (data not shown) to guide annotation transfer to the integrated dataset.

Weighted gene co-expression network analysis (hdWGCNA)^7^ was performed on the integrated dataset using a soft-thresholding power of 12 to approximate scale-free network topology. Gene modules were defined as sets of co-expressed genes with similar expression profiles across cells. Module eigengene matrices were then used as input for single-cell gene regulatory network inference using SCENIC^8^. In SCENIC, transcription factor (TF) regulons were constructed by first identifying target genes with significant expression correlation to a given TF, followed by motif enrichment analysis using the cisBP and RcisTarget databases to confirm the presence of TF binding motifs in the regulatory regions of candidate targets. Regulons were defined as TFs with sets of target genes showing both co-expression and motif enrichment. Gene network analysis was performed using GeneMANIA to predict functional associations among SCENIC-inferred target genes. The resulting interaction networks were visualized and analyzed using Cytoscape (v3.9.1).

### 11. Bioinformatic analysis for bulk analysis

Raw sequencing data were processed using the 3’SRP pipeline, and FASTQ files were aligned to the human reference genome (GRCh38-2020-A). Resulting raw expression matrix was further analysed as described in the 3’SRP pipeline. Counts are then normalized by DESeq2 method. DEGs among condition were determined using DESeq2, and defined as genes with an adjusted p-value < 0.05 and a log2 fold-change > 0.58 (i.e. fold-change > 1.5). WGCNA was performed with a soft threshold of 18^9^. Hierarchical clustering heatmaps were generated by ComplexHeatmap package in R^10^. Gene Ontology enrichment analyses were performed using ClusterProfiler on DEGs for each time points individually^11^. Cell profiles in aggregates were inferred through cross-signature matching using curated gene sets for NC, NP^12–19^, and AF^18,20,21^ cells, as well as for LROs^21–23^ as vacuole markers from published transcriptomic datasets spanning multiple developmental stages and species. Overlap and visualization were performed using ComplexHeatmap^10^ and ggplot2. Fig. S8 summarizes the bioinformatic analysis performed.

Gene set variation analysis (GSVA) ^25^ was performed to obtain a pathway-level representation of transcriptional differences across experimental conditions and time points. Gene sets corresponding to Gene Ontology Biological Process (GO-BP) terms were obtained from the Molecular Signatures Database (MSigDB) using the msigdbr R package. Human GO-BP gene sets were retrieved using the C5 collection and the GO subcollection. Gene sets were converted into a list of gene symbols, with each element corresponding to one GO-BP term. GSVA scores were calculated using the GSVA Bioconductor package (version ≥2.0) using the parameter-based gsvaParam() interface. The variance-stabilized expression matrix and the GO-BP gene sets were provided as input, and a Gaussian kernel (kcdf = “Gaussian”) was used for the expression data. The analysis generated a continuous enrichment score for each GO-BP gene set in each sample, thereby providing a sample-level estimate of the relative activity of each biological process.Gene sets with zero variance across samples were excluded from subsequent analyses. The resulting GSVA matrix was used for downstream statistical analysis and visualization. To assess global differences in pathway activity between experimental groups, a linear model was fitted to the GSVA scores using the limma package. Experimental groups were incorporated into a design matrix without an intercept, and empirical Bayes moderation was applied using the eBayes() function. Global differences across the experimental groups were assessed using the moderated F-test implemented in topTableF(). P-values were adjusted for multiple testing using the Benjamini-Hochberg method, and pathways with an adjusted p-value <0.05 were considered significant. For the focused analysis of extracellular matrix and cartilage-associated programs, selected GO-BP terms related to the biological processes of interest were visualized individually using boxplots, with individual samples displayed as jittered points. These included cartilage morphogenesis, connective tissue development, regulation of extracellular matrix organization, chondroitin sulfate proteoglycan metabolic process, collagen metabolic process, collagen fibril organization, chondroitin sulfate proteoglycan biosynthetic process, and extracellular matrix assembly.

Scripts and list lists can be found here: https://github.com/w-julie/3D-cell-paper. All raw sequencing data will be made publicly available upon publication.

### 12. Visium HD Spatial Gene Expression

For spatial transcriptomics, 5 μm FFPE aggregate sections were processed using the Visium HD Spatial Gene Expression platform according to the manufacturer’s protocol (10x Genomics). Spatial transcriptomics data were with Space Ranger v4.0 to obtain segmented outputs corresponding to polygon-based cellular regions. The Space Ranger output directory containing gene expression matrices, spatial metadata, and segmentation geometries was imported into R (v4.3+) using the Seurat framework. Data were loaded with the Load10X_Spatial() function using multiple bin resolutions, including polygon-level segmentation, and polygon-resolved gene expression data were stored in the Spatial.Polygons assay. Spatial features and segmentation geometries were visualized using SpatialFeaturePlot(), with optional rendering of polygon boundaries and spatial cropping based on coordinate limits for high-resolution inspection of selected tissue regions. The Spatial.Polygons assay was normalized using SCTransform, and the resulting variance-stabilized data were stored in a new assay (Polygon) for downstream analyses. Dimensionality reduction was performed using principal component analysis, and the first 30 principal components were used to construct a shared nearest neighbor graph. Unsupervised clustering was carried out using the Louvain algorithm (resolution = 0.2), followed by UMAP embedding for visualization. Cluster identities were projected back onto tissue space using SpatialDimPlot() to assess spatial organization. Cluster-specific marker genes were identified using a Wilcoxon rank-sum test, retaining only positively enriched genes detected in at least 15% of polygons with a minimum log2 fold-change of 0.25. Mitochondrial and ribosomal genes were excluded prior to downstream analysis. Marker genes were ranked by average log2 fold-change, and the top markers per cluster were used for visualization and data export.

Gene signature activity was quantified at the polygon level using UCell^26^, a rank-based method robust to sequencing depth variation. Curated gene sets used in the 3’SRP were matched to expressed genes and scored using AddModuleScore_UCell(). UCell scores were computed only for polygons with sufficient gene coverage to support rank-based enrichment; polygons with insufficient transcript complexity were not assigned scores. Signature scores were visualized both in low-dimensional embeddings and in spatial coordinates using polygon-resolved spatial plots.

## Data availability

The raw single-cell, bulk and spatial transcriptomics datasets will be deposited into appropriate database.

## Code availability

The code used to generate the figure will be made publicly available on a Github repository.

## Acknowledgments

We thank the GenoA, GenoBIRD, MicroPICell and iPSC core facilities, all supported by IBiSA and Biogenouest, for the use of their resources and technical support. The authors acknowledge the SC3M facility of the Inserm/Nantes Université/ONIRIS UMR1229 RMeS Laboratory and SFR Bonamy. Spatial transcriptomic analyses were supported by the Genomic Technologies Core Facility at the University of Manchester. We acknowledge Boris Halgand and Lisanne T. Laagland for their valuable technical advices. This work was supported by fundings from European Union’s Horizon 2020 research and innovation program (iPSpine; grant agreement number 825925) and the French Society of Rheumatology (SPHERODISC-II), for AC. Funding for CTD was obtained via a Medical Research Council (MRC) award to SMR [MR/W019418/1].

## Author Contributions

Conceptualization, A.C. with advice from L.D., M.A.T., J.W., N.V. and J.G.; Methodology and data curation, A.C., M.B., C.C.^1^, C.C.^4^, C.D., C.M., L.D., R.D., N.V., and J.W.; Project administration, A.C., B.G., J-M.L. and J.G.; formal analysis and programming, B.G.H., and J.W.; writing-original draft preparation, A.C. and J.W.; review and editing of the manuscript, A.C., M.A.T., L.D., S.M.R, N.V., J.W., J-M.L. and B.G.; supervision, A.C. All authors contributed to the interpretation of the data, read and approved the final manuscript.

## Competing interests

The authors declare no competing interests.

Correspondence and requests for materials should be addressed to Anne Camus.

**Figure sup 1. Characterization and cross-validation of cluster identities and MEPC transcriptional signatures.** (a) Distribution of cell clusters across experimental conditions, showing the relative contribution of each condition to the identified cell populations. The two hiPSC lines are indicated independently. The day 7 condition includes day 7 NOTO transfected cells, untreated and treated with a TGF-β inhibitor at different time points. No difference in the transcriptomic analysis was identified. (b) Pearson correlation of pseudo-bulk gene expression profiles across notochord populations stratified by time point and cell line, and compared to an independent dataset showing high degree of correlation. (c) Projection of a murine MEPCs gene signature onto the dataset (Masamsetti et al. 2025^53^), highlighting its enrichment within the MEPCs cluster.

**Figure sup 2. MEPCs and NLCs differentiation efficiency in monolayer cell samples used to generate aggregates**. (a,b) Quantification of TBXT/FOXA2 (a), TBXT/SOX9 (b) immunopositivity of the differentiated cells. Mean values for day 2 (MEPCs), and day 7 NOTO transfected (NLCs) cells are represented in %, for N = 4 biological replicates and n = 3 technical replicates, using 3 distinct hiPSC lines. (c, d) Detailed quantification of the values represented in the bar chart. MEPCs correspond to TBXT^+^/FOXA2^+^ and TBXT^+^, and TBXT^+^/SOX9^+^ and TBXT^+^. NLCs correspond to TBXT^+^/FOXA2^+^ and TBXT^+^, and TBXT^+^/SOX9^+^, TBXT^+^, and SOX9^+^.

**Figure sup 3. Only MEPCs_NPM aggregates are similar to native NP cells.** (a) Heatmap of the expression levels of NP and AF markers using previously published data found in the entire bulk RNA-seq dataset. (b) Heatmap of the expression level of NP markers found in the top 100 genes contributing to PC2. Each row represents one gene and each column a sample. The expression level is shown colored from lowest (blue) to highest (red).

**Figure sup 4. From day 7 and onwards, the MEPCs_NPM aggregates display a transcriptomic signature of matrix-producing cells**. (a-c) GO terms associated with the positively differentially expressed genes at day 7 (a), 14 (b), and 28 (c) in MEPCs_NPM aggregates. (d-e) MAplot showing differential gene expression between day 14 MEPCs_NPM aggregates and all other samples collected at day 14 (a), and between day 28 MEPCs_NPM aggregates and all other samples collected at day 28 (b). Each point represents a gene, plotted according to its mean expression level across all samples from the same timepoint (average log expression; x-axis) and its log2 fold change in MEPCs_NPM relative to other conditions (y-axis). Genes with a mean expression value greater than 1 on the x-axis are highlighted, emphasizing transcriptional changes among robustly expressed genes. Significantly upregulated and downregulated genes (adjusted p ≤ 0.05) are indicated in color, while non-significant genes are shown in grey.

**Figure sup 5. Chondrogenic differentiation induces the emergence of off-target cell gene signatures, while in the NPM conditions the specific transcriptomic signature is preserved.** (a-d) Identification of neural related differentiation in the chondrogenic differentiation, with differentially expressed genes shown in the MAplots at day 14 (a) and day 28 (c), and their associated GO terms (b, d). (e) Identification of major WGCNA modules of co-expressed genes distinguishing global transcriptomic profile depending on culture conditions. Each row represents a WGCNA module and each column a sample. The number of genes in the module is indicated. The expression is shown colored from lowest (blue) to highest (red)

**Figure sup 6. NLCs aggregate matrisome is distinct from native NP cells in all culture conditions.** Temporal expression profiles of key genes from the WGCNA-identified module across differentiation (days 7, 14, and 28). Line plots display centered expression dynamics in NLCs_CHM aggregates (left), CHM-NPM (middle), and NPM (right). Genes are grouped by functional categories: core matrisome (collagens, glycoproteins), ECM-affiliated proteins, matrisome-associated proteins, and growth factor-related factors.

**Figure sup 7. Immunohistochemistry analysis of NP markers in NLCs_NPM aggregates from day 14 to day 28.** (a) Violin plots showing normalized gene counts from bulk RNA-seq for NP markers (*SOX9*, *KRT8*, *CD109*, *CD166, Tie2*). Each dot represents an individual sample, shape-coded by cell source. (b) Representative immunohistochemistry for NP markers at day 14, 21 and 28 in NLCs_NPM aggregates. Scale bars, 50 µm; insets, 25 µm.

**Figure sup 8. Schematic workflow of bioinformatic analysis.** (a) for the scRNAseq data. (b) for the bulk RNAseq data. (c) for the spatial transcriptomics.

**Table S1. Differentially expressed genes used to annotate single-cell clusters**

**Table S2. Top100 genes contributing to PC1 to PC4.** * Indicates genes that are found in the curated NP gene list from published transcriptomic datasets spanning multiple developmental stages and species. ** indicates genes related to chondrocyte identity.

**Table S3. Summary of antibody incubation conditions used for immunostaining**

